# Oral administration of an anti-CfaE secretory IgA antibody protects against Enterotoxigenic *Escherichia coli* diarrheal disease in a non-human primate model

**DOI:** 10.1101/748442

**Authors:** Matteo Stoppato, Carlos Gaspar, James Regeimbal, Gladys Nunez, Serena Giuntini, Melissa A. Gawron, Jessica R. Pondish, Joseph C. Martin, Matthew Schneider, Zachary A. Schiller, Mark S. Klempner, Lisa A. Cavacini, Yang Wang

## Abstract

Enterotoxigenic *Escherichia coli* (ETEC) is a leading cause of diarrhea-associated illness in developing countries. There is currently no vaccine licensed to prevent ETEC and the development of an efficacious prophylaxis would provide an intervention with significant impact. Recent studies suggested that effective protection could be achieved by inducing immunity to block colonization of ETEC. Here, we evaluated the efficacy of secretory (s) IgA2 and dimeric (d) IgA2 of an anti-colonization factor antigen antibody, 68-61, in the *Aotus nancymaae* non-human primate (NHP) ETEC challenge model via oral and parental delivery. Thirty-nine animals were distributed across 3 groups of 13, and challenged with 5.0×10^11^ cfu of H10407 on Day 0. Group 1 received a dIgA2 68-61 subcutaneously on day 0. Group 2 received a SIgA2 68-61 orally on days −1, 0, and +1, and Group 3 received an irrelevant SIgA2 antibody orally on days −1, 0, and +1. All animals were observed for symptoms of diarrhea, and stools were collected for ETEC colony counts. SIgA2 treatment significantly lowered the attack rate, resulting in a protective efficacy of 71.4% (p=0.025) in Group 2 as compared to Group 3. Anti-CfaE dIgA2 treatment group reduced the diarrheal attack rate, although the reduction did not reach significance (57.1%; P=0.072) as compared to the irrelevant SIgA2 Group 3. Our results demonstrated the feasibility of oral administration of SIgA as a potential immunoprophylaxis against enteric infections. To our knowledge, this is the first study to demonstrate the efficacy of administrated SIgA in a non-human primate model.

## INTRODUCTION

Enterotoxigenic *Escherichia coli* (ETEC) is the most common cause of diarrheal illness in infants in the developing world and in travelers to endemic countries. An estimated 10 million cases per year occur among travelers and military personnel deployed in endemic regions [1]. ETEC is a non-invasive pathogen that mediates small intestine adherence through filamentous bacterial surface structures known as colonization factors (CF). Once bound to the small intestine, the bacteria produce toxins causing a net flow of water from enterocytes, leading to watery diarrhea [2, 3]. Previous approaches to prevent ETEC infection have targeted bacterial attachment and colonization. However, poor responses to vaccines and difficulties in the establishment of protective mucosal immunity against diverse types of CFs have hindered the licensing of ETEC vaccines.

CFA/I is one of the most prevalent CFs expressed by pathogenic ETEC strains. CFA/I is composed of a stalk consisting of a long homopolymeric subunit and a minor adhesin subunit (CfaE) at the tip of the fimbria. Recent studies have demonstrated that the adhesin subunit can induce anti-adhesive immunity against ETEC infection [4, 5]. In human immunoprotection trials, oral administration of anti-CfaE bovine IgG provided protection in over 60% of the test group, suggesting that an adhesin-based vaccine could be effective to elicit endogenous production of protective antibodies [6].

IgG and secretory IgA (SIgA) are both present in the small intestine as effector molecules of *mucosal* immune system. SIgA is considered to be one of the most important effector molecules because it constitutes the primary immune defense against pathogens at the mucosal surface [7]. In secretory IgA, two IgA monomers are covalently linked by a joining chain (J-chain), and stabilized by a polypeptide called the secretory component that make the molecule more resistant to digestion in the small intestine than IgG [8]. Early studies also have suggested that the secretory component may have its own antimicrobial activity to block epithelial adhesion of ETEC [9].

Our laboratory has recently identified a panel of anti-CfaE human monoclonal antibodies IgGs that could be employed as an immunoprophylaxis to prevent ETEC diarrhea [10]. We performed isotype switch to SIgAs and we are investigating the potential of these SIgAs to serve as immunoprophylaxis against ETEC diarrhea. Here, we evaluated the efficacy of one lead anti-CfaE SIgA, 68-61, in the *A. nancymaae* non-human primate ETEC challenge model (ETEC strain H10407) [11, 12]. In this model, *A. nancymaae* has been shown to be susceptible to diarrhea in response to experimental infection with ETEC expressing CFA/I, mimicking ETEC pathogenesis in human (Jones 2006). 68-61 was administered to *Aotus* either as a dimeric IgA2 (dIgA2) via a single subcutaneous dose (SC) on the day of challenge (day 0) or as a secretory IgA2 (SIgA2) via oral delivery on days −1, 0 (challenge day), and +1. Animals were then monitored for diarrhea as previously described [12]. Our results demonstrate that oral administration of SIgA2 can protect animals from diarrhea associated with ETEC infection.

## RESULTS

### Production and characterization of 68-61 SIgA2 and dIgA2 antibodies for NHP studies

Large scale production for anti-CfaE 68-61 dIgA2 and SIgA2 antibodies were set up to generate sufficient material for NHP studies using an established IgA production platform in our laboratory [10]. To verify the antibody quality, purified antibodies were analyzed by SDS-PAGE and western blots (Fig. 1A). MRHA and Caco-2 cell adhesion assays were also conducted to test the antibody in vitro functionalities. Similar to what was reported previously [10], both purified 68-61 dIgA2 and SIgA2 showed functional activity in both hemagglutination assay (minimal inhibitory concentration of 0.04ug/ml and 0.08ug/ml) and Caco-2 adhesion assay (Fig. 1B and C, respectively).

**Figure 1:**
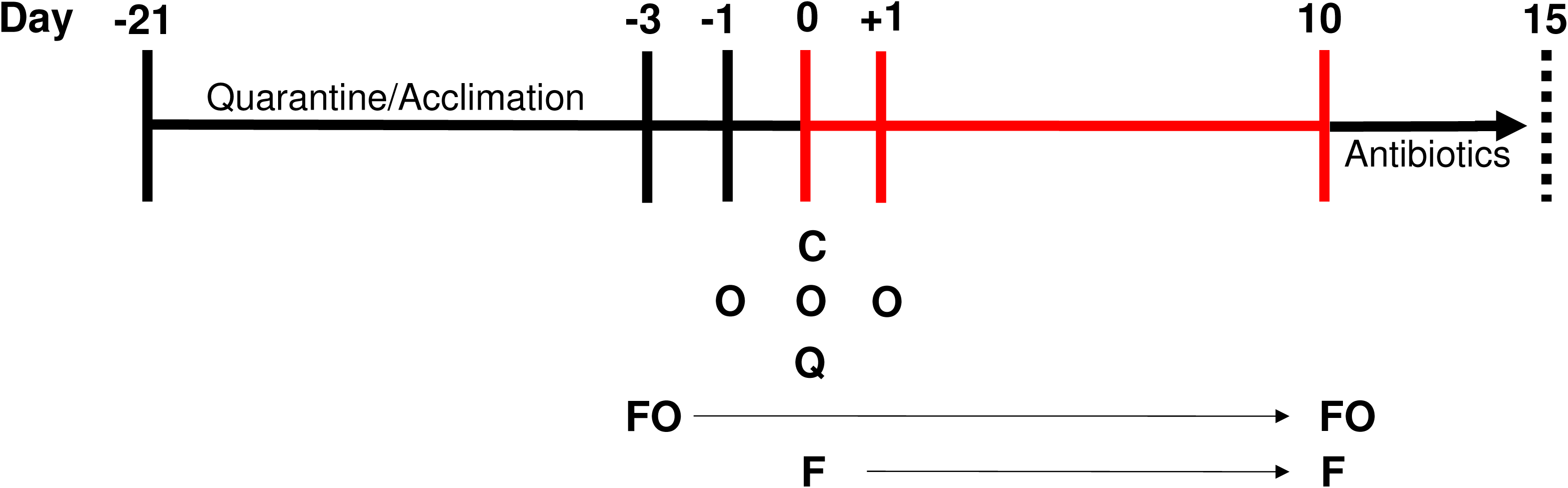
Study design of the challenge experiment. FO = fecal observation and scoring, all animals; F = fecal collection, all animals; C = challenge; O = oral antibody administration; Q= SubQ antibody administration.

**Figure 2:**
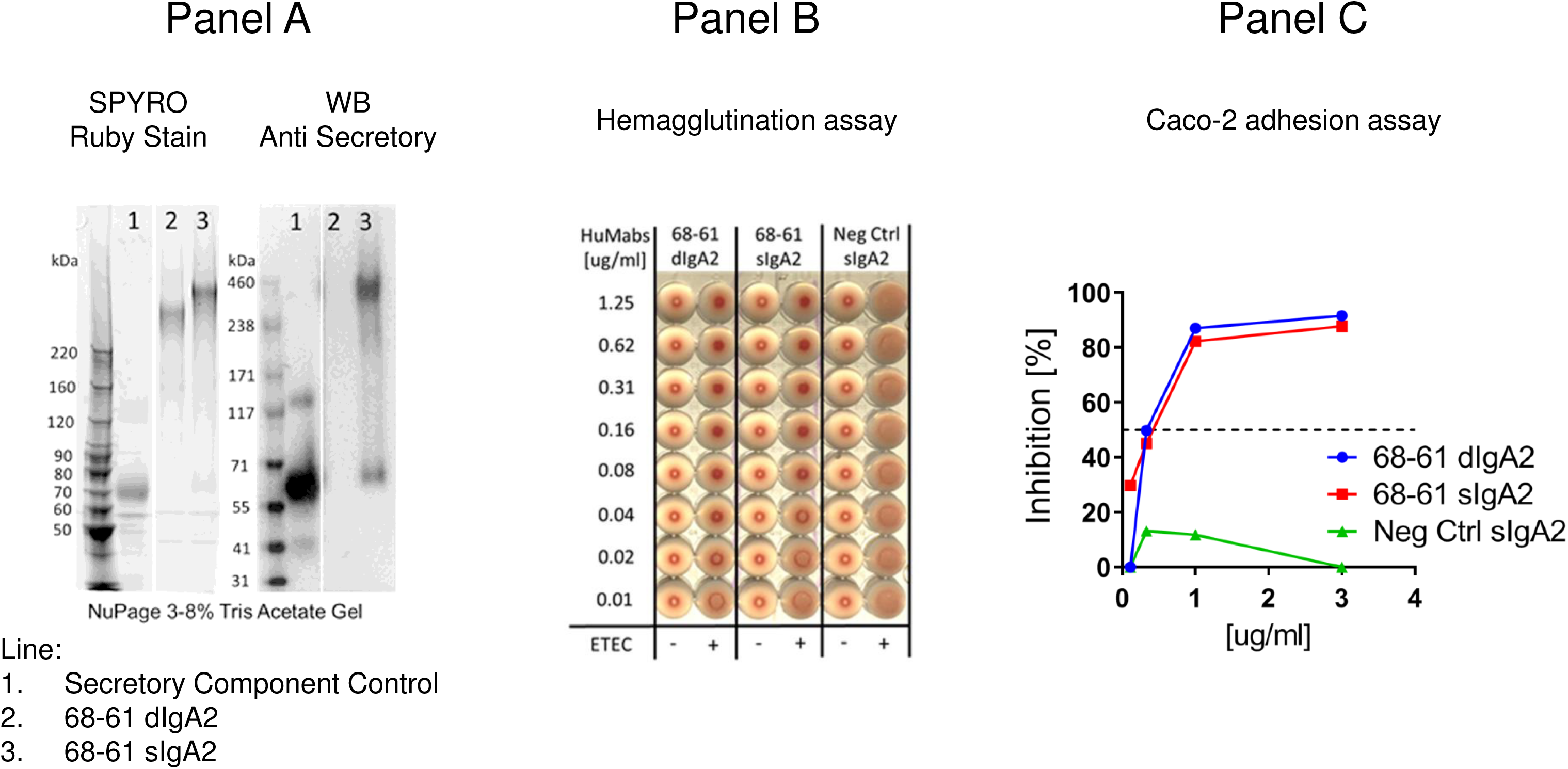
Characterization of 68-61 dimeric and secretory IgA. A) SDS-PAGE and Western blot of 68-61 dIgA (line 2) and 68-61 SIgA (line 3). Antibody specific to secretory component and SIgA was used for western blot. B) Activity of 68-61 dIgA2 and SIgA2 in a mannose resistant hemagglutination assay of human group A erythrocytes. The minimal inhibitory concentration to prevent hemagglutination is 0.04 and 0.08 μg/ml for dIgA2 and SIgA2 respectively. The assay was repeated three times using different blood donors. C) Functionality of 68-61 dIgA2 and SIgA2 tested in a representative Caco-2 adhesion assay

### Antibody efficacy study in a non-human primate model challenged with ETEC

Dimeric (Group 1) and secretory (Group 2) anti-CfaE IgA2 antibodies were administered to *Aotus nancymae* monkeys to determine their efficacy against ETEC H10407 strain. Animals administered the irrelevant control SIgA2 antibody (Group 3, oral) had a diarrheal attack rate of 58% (7/12), within the range of the reported attack rate in this animal model [12]. Anti-CfaE dIgA2 treatment (Group 1; S.C.) lowered the attack rate to 23% (3/13), while SIgA2 treatment (Group 2; oral) significantly lowered the attack rate to 15% (2/13) as compared to Group 3. One animal in Group 3 was excluded from analysis due to the onset of diarrhea prior to challenge (Table 2). There was no significant difference in the colonization rate or the duration of shedding between the treatment groups. Based on the diarrheal attack rates, oral anti-CfaE SIgA2 (Group 2) treatment resulted in a protective efficacy of 71.4% (P=0.025) compared to the irrelevant SIgA2 (Group 3). Treatment with a subcutaneous injection of anti-CfaE dIgA2 (Group 1) reduced the diarrheal attack rate, although the reduction did not reach significance (57.1%; P=0.072) as compared to Group 3 (Table 3). Of note, Group 1 animals did not receive any of the oral rehydration drink on days −1 and +1 that was used to orally administer the SIgA antibodies in Groups 2 and 3.

**Table 1.**
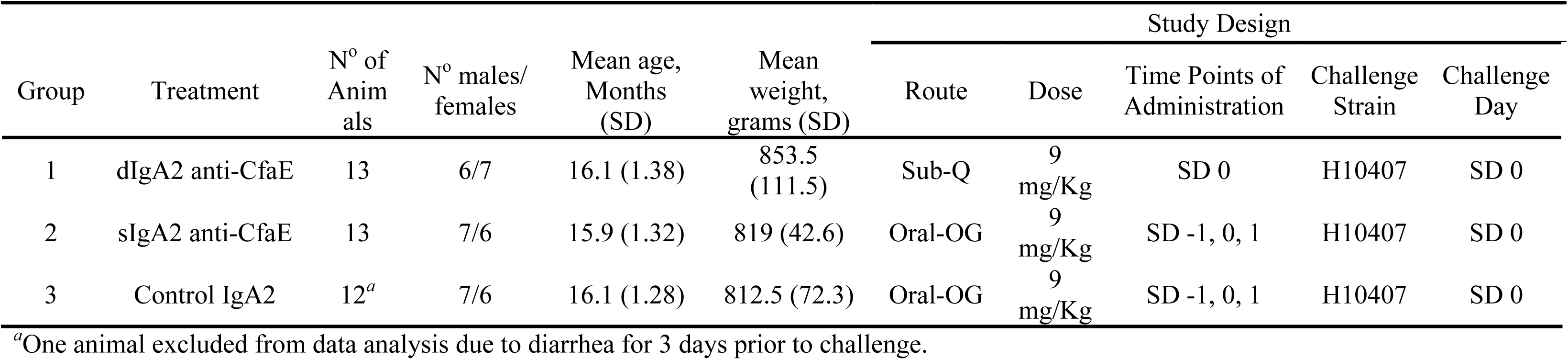
Demographic variable and study design

**Table 2.**
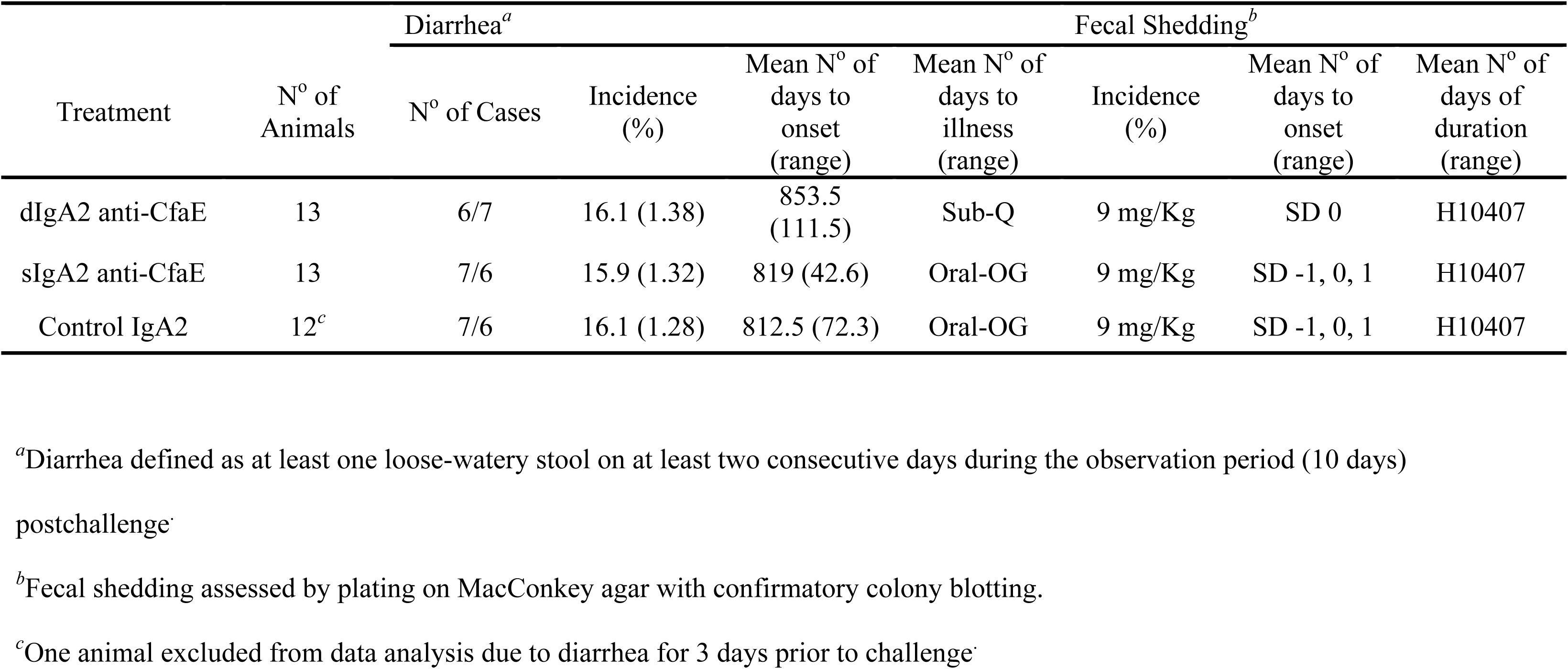
Diarrhea and colonization after oral challenge of *A. nancymaae* with ETEC strain H10407

**Table 3.** Protective Efficacy in *A. nancymaae*

| Group | Vaccine | N° with diarrhea/N (%) | Protective Efficacy (%) | *P* value<sup>a</sup> |
| --- | --- | --- | --- | --- |
| 1 | Dimeric anti-CfaE IgA2 | 3/13 (23) | 57.1 | 0.072 |
| 2 | Secretory anti-CfaE IgA2 | 2/13 (15) | 71.4 | 0.025 |
| 3 | Control IgA2 | 7/12 (58) | - | - |

<sup>a</sup>Fisher Exact Test, two-tailed, comparing frequency of diarrhea test groups to control group.

## DISCUSSION

While both IgG and IgA are expressed at the mucosa, IgA is usually more effective on a molar basis and thus are the natural choice for mucosal passive immunization. The avidity of mucosal IgA, due to multimeric structure, enhances antibody binding with antigens and increases antibody mediated conformational or structural changes in the antigen. The diverse, high level of glycosylation of IgA antibodies, in comparison to IgG, further protects the mucosal surface by non-specific interference with microbial adherence. Here, we further explored the feasibility of administration of anti-CfaE IgA for protection against an ETEC challenge in the *A. nancymaae* non-human primate model. Oral delivery of an anti-CfaE SIgA2 resulted in 71.4% protective efficacy against ETEC diarrhea in animals.

The administration of a single dose of anti-CfaE dIgA antibody subcutaneously resulted in a 57.1% reduction of ETEC diarrhea. Though not significant (P=0.072), in contrast to the control animals (Group 3), Group 1 animals did not receive any of the oral rehydration drink, which may have imparted a small therapeutic effect to the control animals not recapitulated in Group 1. Eliminating this difference, dose optimization, and/or temporal administration experiments may reveal a significant reduction in diarrhea and further experimentation is clearly warranted.

Nevertheless, fecal shedding of H10407 was observed in all animals regardless of the treatment (Table 2). These results are consistent with observations in previous vaccine studies in animals and human challenge volunteers [12, 13], suggesting that the anti-ETEC immunity may inhibit bacterial adherence and/or pathogenicity without affecting fecal shedding. Histological analysis in future experiments may verify the prevention of intestine colonization and better define the protective mechanism. Regardless, these results are promising and further definition of dose and temporal kinetics of SIgA oral administration should further increase efficacy.

IgA antibodies function in mucosal immunity as the first line of defense against pathogens making them attractive as candidate therapeutics [14]. However, efficacy studies of passively administered IgA in animal models has been limited by the capabilities of large-scale production in the laboratory. In this study, we were able to utilize our recently developed IgA production methods [10] to generate high quality dIgA and SIgA to support a well-powered non-human primate animal study. Through our study, we established the feasibility of oral administration of SIgA2 as a potent immunoprophylaxis against enteric infections. We further demonstrate the potential feasibility of subcutaneous administered dIgA2 in preventing ETEC diarrhea. These results are of great interest as they demonstrate for the first time that SIgA2 can be used as a preventative against bacterial diarrhea.

## MATERIALS AND METHODS

### 68-61 SIgA and dIgA antibody production and characterization

68-61 dIgA2 and SIgA2 antibodies were produced and characterized as previously described [10]. To generate dIgA2, the heavy and light chain vectors were co-transfected with a J chain expressing vector with equal molar ratio in CHO cells. For SIgA2, a secretory component expressing vector was added to the dIgA2 transfection reaction (equal molar ratio for all vectors). Supernatant was run through a column of Capture Select IgA (ThermoFisher) or CaptoL resin (GE Life Sciences) for dIgA and SIgA respectively, followed by size exclusion chromatography (HiLoad 26/600 Superdex 200 pg size exclusion column; GE Life Sciences) to separate out the desired dimeric or secretory antibodies. Desired fractions were pooled, concentrated and quality tested by SDS-PAGE, western blots and through mannose-resistant hemagglutination (MRHA) and Caco-2 cell adhesion assays.

### Mannose resistant hemagglutination assay of human group A erythrocytes

ETEC cultures were taken from frozen cell banks and diluted in sterile 0.15 M saline solution until reaching an OD_600nm_ of 1 for the assay. Human erythrocytes type A+ stored in K3EDTA were washed three times with 0.15 M saline solution and resuspended in the same solution to a final concentration of 1.5% (vol/vol). In a U-bottom 96-well plate (Nunc Thermo Scientific) 100 µl of IgA antibodies were added in duplicate to the top row and diluted 1:2 down the plate in 0.15 M saline solution. 50 µl of appropriately diluted ETEC was added to each well together with 50 µl of 0.1 M D-mannose solution (sigma). The plate was incubated for 10 minutes at room temperature. After incubation, 50 µl of blood solution was added to the plate and mixed well (200 µl final volume). Plates were allowed to sit stagnant at 4°C for two hours.

Hemagglutination was then observed without the aid of magnification. The absence of a pellet of red blood cells at the bottom of the well is indicative of positive hemagglutination. Blood was ordered fresh every week (BioreclamationIVT).

### Caco-2 adhesion assay

Caco-2 cells seeded at 1 × 10^5^ cells/mL were grown in 24-well tissue culture plates containing Dulbecco’s modified Eagle’s medium (DMEM), at 37°C in 5% CO_2_ static. Frozen bacterial banks were streaked on CFA agar plates and grown overnight at 37°C. The next day, bacteria were resuspended in PBS and diluted until reaching an OD_600nm_ of 0.1. Antibody dilutions were set up in a deep well plate. Antibody dilutions and bacteria were combined in a 1:10 ratio and allowed to shake at 300 rpm for one hour at room temperature. After incubation, 0.2 mL of antibody/bacteria mixture was added to each well containing Caco-2 cells. The cells were then incubated statically for 3 hours at 37°C. Cells were then washed four times with 1 mL PBS to remove non-adherent ETEC cells. Afterwards, Caco-2 cells were dislodged with 0.2 mL 0.25% trypsin. Cells were collected via gentle centrifugation and resuspended in 1mL PBS. Dilutions were plated on CFA agar plates and colonies counted the next day. IC_50_ was defined as concentration of antibody needed to inhibit 50% of ETEC adhesion to the Caco-2 cells, compared to an irrelevant isotype antibody.

### Ethics statement

The non-human primate (NHP) research was conducted in an AAALAC-accredited Laboratory Animal Facility, in compliance with the Animal Welfare Act and in accordance with principles set forth in the “Guide for the Care and Use of Laboratory Animals,” Institute of Laboratory Animals Resources, National Research Council, National Academy Press, 1996, and other U.S. Federal Government statutes. Local approval was by the U.S. Naval Medical Research Unit No 6 (NAMRU-6) Institutional Animal Care and Use Committee (IACUC), second-level approval from the U.S. Navy Bureau of Medicine and Surgery, and the study was approved via Resolucion Directoral No. 0452-2017 SERFOR-DGGSPFFS by the Forestry and Wild Fauna Service, Peruvian Ministry of Agriculture

The *A. nancymaae* used in this study were purchased from the Instituto Veterinario de Investigaciones Tropicales y de Altura (IVITA), University of San Marcos, Lima, Peru, and shipped to NAMRU-6 in Lima, Peru. Animals were maintained in pairs when not required to be individually housed for sample collection, fed a standard monkey diet supplemented with fruit and provided water *ad libitum*.. Following the study, antibiotic treated and ETEC-free animals were returned to the IVITA colony.

### Administration of antibody and ETEC challenge inoculums

The ETEC challenge model has been previously described [10, 12]. Briefly, *A. nancymaae* monkeys were screened by enzyme-linked immunosorbent assay (ELISA). Animals deemed seropositive were excluded from the study. The remaining thirty-nine animals were distributed across three groups of 13 according to age, sex, and weight. Following a 21 day acclimation period, the animals were fasted overnight and on study day 0 all animals were anesthetized with ketamine hydrochloride (10mg/kg weight, Ketalar, Parke-Davis) and an orogastric feeding tube was placed (5Fr/Ch, 1.7 mm X 41 cm). All animals also received ranitidine (1.5 mg/kg) by intramuscular injection 90 minutes prior to challenge to inhibit gastric acid production, and 5 ml CeraVacxII (CeraProducts, Jessup, MD) was given 30 minutes prior to challenge to neutralize stomach pH. All animals were then challenged with 5×10^11^ cfu ETEC CFA/I+H10407 (5 ml volume).

All groups received an antibody treatment (9 mg/kg) prior to challenge on Day 0. Group 1 received an anti-CfaE dIgA2 antibody by subcutaneous (S.C.) injection. Group 2 received an anti-CfaE SIgA2 antibody via the orogastric line. Group 3 received a control SIgA2 antibody against an HIV target (no cross-reactivity with H10407 in vitro, data not shown) via the orogastric line. Group 2 and Group 3 also received antibody treatment one day prior to challenge (day -1) and one day post challenge (day +1). These additional treatments (9 mg/kg) were prepared by diluting the antibody (anti-CfaE SIgA2 for Group 2 and control SIgA2 for Group 3) into 5 mL total volume of Prang oral rehydration drink (Bio-Serv; orange flavor), and the diluted antibody was then administered orally by syringe via voluntary consumption. All animals were observed for 10 days and then treated with Enrofloxacin until ETEC H10407 was not detected in stool samples. The Study design of the challenge model is illustrated in Figure 1. The demographic variables of animals in each individual group is listed in Table 1.

### Observation after passive immunoprophylaxis and challenge

Animals were observed twice daily for signs and symptoms of diarrhea starting on study day -3 and continuing for 10 days after challenge. Stools were graded as follows: grade 1 (formed, firm stool pellets), grade 2 (formed but soft stool pellets or droppings), grade 3 (loose, unformed feces), grade 4 (watery, non-clear feces), and grade 5 (watery, clear liquid stools). Stools graded 1 or 2 were considered normal, whereas stools graded 3, 4, or 5 were considered abnormal. The case definition of a diarrhea episode was defined as the passing of grade 3 or higher stools for at least two consecutive days during the observation period. The duration of diarrhea was defined as the time between the first day of a diarrhea episode and the last day of diarrhea preceding two consecutive diarrhea-free days. Animals meeting the case definition of diarrhea prior to the challenge were excluded from data analysis.

Fecal cultures for H10407 ETEC were performed daily for 10 days after challenge by streaking fresh stool and plating serial dilutions of stool directly onto MacConkey agar. Presumptive H10407 isolates (lactose-positive) were confirmed by colony blot using rabbit antisera against CFA/I. Stool was considered negative for H10407 if no lactose positive *E. coli* colonies were isolated, or if 10 presumptive colonies were negative by immunoblot. A period of fecal shedding was defined as isolation of H10407 (CFA/I positive colonies) from stool collected after challenge, beginning (onset) as early as the first day after challenge, and ending (duration) on the last day that H10407 is detectable in stool, up to day 10 post challenge.

### Statistical analyses

Intergroup comparison of clinical outcomes were performed using nonparametric tests for continuous outcomes (Kruskal-Wallis test for comparing the values for more than two groups) and Fisher’s exact test for nominal outcomes. All statistical tests were interpreted in a two-tailed fashion with acceptance of significance set at the *P* < 0.05 level.

## ACKNOWLEDGMENTS

The authors declare that there are no financial, institutional or other relationships that might lead to bias or a conflict of interest. The views expressed in this article are those of the authors and do not necessarily reflect the official policy or position of the Department of the Navy, Department of Defense, nor the U.S. Government.

This work was supported by the Defense Advanced Research Project Agency (DARPA-BAA-13-03) to MK.

Some of the authors are military service members or employees of the U.S. Government. This work was prepared as part of their official duties. Title 17 U.S.C. § 105 provides that ‘Copyright protection under this title is not available for any work of the United States Government’. Title 17 U.S.C. § 101 defines a U.S. Government work as a work prepared by a military service member or employee of the U.S. Government as part of that person’s official duties.

We would like to thank the veterinary staff of Naval Medical Research Unit 6 for their technical support, LTC Robin Burke for input regarding health and safety parameters, Ms. Nereyda Espinoza, Ms. Monica Nieto, Mr. Jesús Rojas and Ms. Rosa Castillo for their excellent technical assistance during this study.

